# Age-related differences in performance and fatigability during an isometric quadriceps intermittent fatigue test

**DOI:** 10.1101/2021.09.19.460938

**Authors:** Giorgio Varesco, Eric Luneau, Léonard Féasson, Thomas Lapole, Vianney Rozand

## Abstract

The aim of the present study was to investigate age-related differences in fatigability induced by an isometric quadriceps intermittent fatiguing test in young (<35 years old), old (>60 years old) and very old (>80 years old) men and women. Maximal force loss, contractile function and voluntary activation of the knee extensors were evaluated throughout an isometric fatiguing test using femoral nerve magnetic stimulations. Older adults performed more contractions (index of relative performance) than young (P = 0.046) and very old adults (P = 0.007), without differences between young and very old adults. Total work (absolute performance) was greater for young and old adults compared to very old adults (P < 0.001), without differences between young and old adults. At exhaustion, force loss was greater for young (−28 ± 9%) compared to old adults (−19 ± 8%), but not very old adults (−23 ± 8%). The response to femoral nerve stimulation decreased similarly at exhaustion for the three age groups, indicating similar alteration in contractile function with age. No impairment in voluntary activation was observed. Impairments in neuromuscular parameters were similar for men and women. This study showed that older adults were less fatigable than young adults during an isometric intermittent fatiguing task of the knee extensors. This greater fatigue resistance was not maintained in very old adults independent of sex. Fatigability at exhaustion was likely due to impairments in contractile function for the three age groups.

## Introduction

Healthy ageing is usually accompanied by impairments in neuromuscular function and fatigability, leading to increased risk of autonomy loss and incidence of frailty [1]. These impairments are not linear with age. For example, the decline of maximal isometric force accelerates in very old adults (i.e. >80 years) compared to old adults (i.e. >60 years) [2,3], with greater impairments in women compared to men [4,5]. In a recent meta-analysis, Krüger et al. [6] reported an overall lower performance fatigability (exercise-induced maximal force loss) in older compared to young adults after isometric fatiguing tasks. However, these results are equivocal, with some studies reporting a large effect size (Hedges’s *g* >2) and others reporting no age-related difference or greater fatigability in older adults [6]. Importantly, studies investigating performance fatigability after isometric tasks in very old adults are limited. Allman and Rice [7] compared performance fatigability after an intermittent isometric fatiguing task of the elbow flexors between young and very old adults, and reported similar values between the two age groups. However, the absence of a group of older adults did not allow the rate of decline in neuromuscular function between old and very old adults to be determined. Justice et al. [8] showed that time to exhaustion during an isometric sustained fatiguing task of the ankle dorsiflexors was longer for older adults (65-75 yr) compared to young adults (<35 yr), with similar performance between young and very old adults (75-90 yr). However, performance fatigability was not investigated in this study, nor the mechanisms explaining these age-related differences, *i*.*e*. above (voluntary activation) or below (contractile function) the neuromuscular junction.

In old adults, impairments in voluntary activation following single-limb exercises seem to be similar compared to young adults [6]. Conversely, impairments in contractile function with fatigue show inconsistent results, with similar or lower alterations in old compared to young adults after isometric fatiguing tasks [6]. Again, evidence for alterations in voluntary activation and contractile function in very old adults are scarce. Sundberg et al. [9] observed greater impairments in contractile function but not voluntary activation compared to young and old adults. Because variability increases with advancing age, standardized and validated testing procedures as well as assessments of physical activity levels are needed when comparing groups of different ages [10].

Thus, the present study aimed to investigate performance fatigability of the knee extensor (KE) muscles induced by the isometric quadriceps intermittent fatiguing (QIF) test in young, old and very old men and women. We expected similar performance fatigability in young and very old adults after the isometric intermittent fatiguing test, while older adults would show a lower performance fatigability. We further hypothesized lower impairments in contractile function in old adults than very old and young adults, without age-related differences in voluntary activation impairments.

## Methods

### Participants

Thirty young adults [15 men (YM); 15 women (YW)], 19 old adults [10 men (OM); 9 women (OW)] and 30 very old adults [15 men (VOM); 15 women (VOW)] participated in the study and were clearly informed on the experimental procedures prior giving their written consent. Participants were free from psychological, musculoskeletal and neurological disorders or disability, and were asked to avoid exercise and alcohol consumption for 24 h before each visit. All the procedures were approved by the Comité de Protection des Personnes and were performed in accordance with the declaration of Helsinki (2013; ClinicalTrials.gov identifier: NCT02675192). The University Hospital of Saint-Etienne (France) was the sponsor of this study.

### Experimental protocol

Participants visited the laboratory on two separate occasions. During the first visit, all participants were familiarized with the neuromuscular testing procedures, and an accelerometer (wGT3X-BT; ActiGraph, Pensacola, USA) was given to wear for 7 consecutive days to objectively assess physical activity [11]. The old and very old adults also performed a 6-minutes-walk-test (6MWT), which consisted in covering the longest distance possible in 6 minutes in a corridor of 30 meters [12]. One week later during the second visit, KE function was evaluated using the isometric QIF test [13].

### Experimental setup

Participants were seated upright on a custom-build chair with the hip angle set at 120° (180°= full extension) to facilitate coil placement over the femoral triangle (see below). Knee angle was fixed at 90°. The right leg was systematically evaluated other than for specific reasons (i.e. operated knee, unilateral pain). In these instances, the left leg was tested (1 OW, 2 VOM, 1 VOW). Extraneous movements of the upper body were minimized using belts across the waist and the thorax. Force of the KE was assessed using a force transducer (Omega Engineering Inc., Stamford, CT, USA) firmly attached over the malleoli, and digitized at a sampling rate of 2 kHz by PowerLab System (16/30-ML880/P, AD Instruments, Bella Vista, Australia).

### Femoral nerve magnetic stimulation

Femoral nerve magnetic stimulation (FNMS) was performed with a 45-mm figure-of-eight coil powered by two linked Magstim 200 stimulators (peak magnetic field: 2.5 T, stimulation duration: 0.1 ms; Magstim, Whitland, UK). The coil was positioned above the femoral triangle to recruit the femoral nerve and all stimuli were given at 100% of the maximum stimulator output. Optimal stimulation site evoking the highest twitch force amplitude was determined with minor adjustments and marked on the skin. FNMS supramaximality was checked with decreasing stimulator power output (100%, 95%, and 90%). Because FNMS effectiveness in deliver supramaximal stimuli is altered when fat thickness under the coil increases [14], clear plateaus in maximal twitch amplitude were not observed for all participants. Thus, only valid responses to FNMS were retained for analysis (YM: 15; OM: 8; VOM: 10; YW: 15; OW: 9; VOW: 8).

### Isometric QIF test

The session started with a warm-up consisting of three ∼3-s contractions at 30%, 50% and 70% of the perceived maximal force. Successively, two 5-s maximal voluntary isometric contractions (MVICs) were performed, separated by 1-min of rest. A third MVIC was performed if the difference between the peak force values was >5%. Participants were instructed to push “as hard as possible” and strong verbal encouragement was provided. Visual feedback of the force signal was displayed on a screen positioned in front of the participants. Then, a neuromuscular evaluation (NME) was performed. NME consisted of a 5-s MVIC during which a 100-Hz doublet (Db_100,s_) was superimposed through FNMS, and a resting potentiated 100-Hz doublet (Db_100_) delivered ∼2 s after relaxation. Once the baseline measures were performed, the QIF test started after at least 1 min of recovery.

The QIF test consisted of incremental sets of 10 intermittent (5-s on / 5-s off) isometric contractions, starting at 10% MVIC for the first set, with increments of 10% MVIC per set until exhaustion [13]. Visual feedback of the target force level was provided, and participants followed a soundtrack with the contraction-relaxation rhythm. A line of 4-N thickness indicated the target force (2 N below and 2 N above). Exhaustion was defined as two consecutive contractions where force decreased below the target force level for 2.5 s. NME was performed immediately (∼2 s) at the end of each set and at task failure. The duration between the end of a set and the beginning of the following one was standardized at 25 s.

### Data analysis

Physical activity data were analyzed using ActiLife software (v6.13.4, ActiGraph, Pensacola, USA) and quantified as steps·day^-1^. All the remaining data were analyzed using Labchart software (v8, ADInstruments). MVIC was calculated as the average of the highest 0.5-s window of the contraction. Performance fatigability was defined as the percentage of MVIC loss from baseline. Total work was calculated as the sum of the force–time integral of each contraction and indicated absolute performance at the QIF test. Total number of contractions performed indicated relative performance. Voluntary activation (VA) was calculated from the formula: VA% = Db_100,s_/Db_100_ × 100. Impairments in contractile function were quantified as the change in Db_100_ from baseline.

### Statistical Analysis

All variables are reported as mean ± SD in the text, tables and figures. Generalized Linear Models were used to analyze the age- and sex-related differences in anthropometric (age, weight, height) and neuromuscular characteristics at baseline (absolute and normalized MVIC, Db_100_, VA), and performance (total work, number of contractions). Generalized Estimating Equations analyses using an autoregressive (AR-1) structure were performed on data expressed relative to baseline to investigate the effects of age and sex across the QIF test on MVIC, Db_100_ and VA [stage (stages 1, 2, 3, 4 and exhaustion) × age (young: YA *vs*. OA *vs*. VOA) × sex (men, women)]. Because participants performed different numbers of stages, data were compared until the last stage performed by all participants (i.e. 4^th^ stage). For all analyses, when significant effects were observed, Bonferroni correction was applied to post-hoc analyses. Significance was set at P ≤ 0.05. Statistical analysis was conducted using SPSS software (Version 23.0, IBM, Chicago, IL).

## Results

### Baseline measurements

Table 1 summarizes the participants’ characteristics. Young and old adults walked significantly more steps·day^-1^ than very old adults (age effect, P < 0.001), with women being more active than men (sex effect, P = 0.013). Very old adults performed shorter distance at the 6MWT than old adults (age effect, P < 0.001) with women performing shorter distance than men (sex effect, P < 0.001).

**Table 1.**
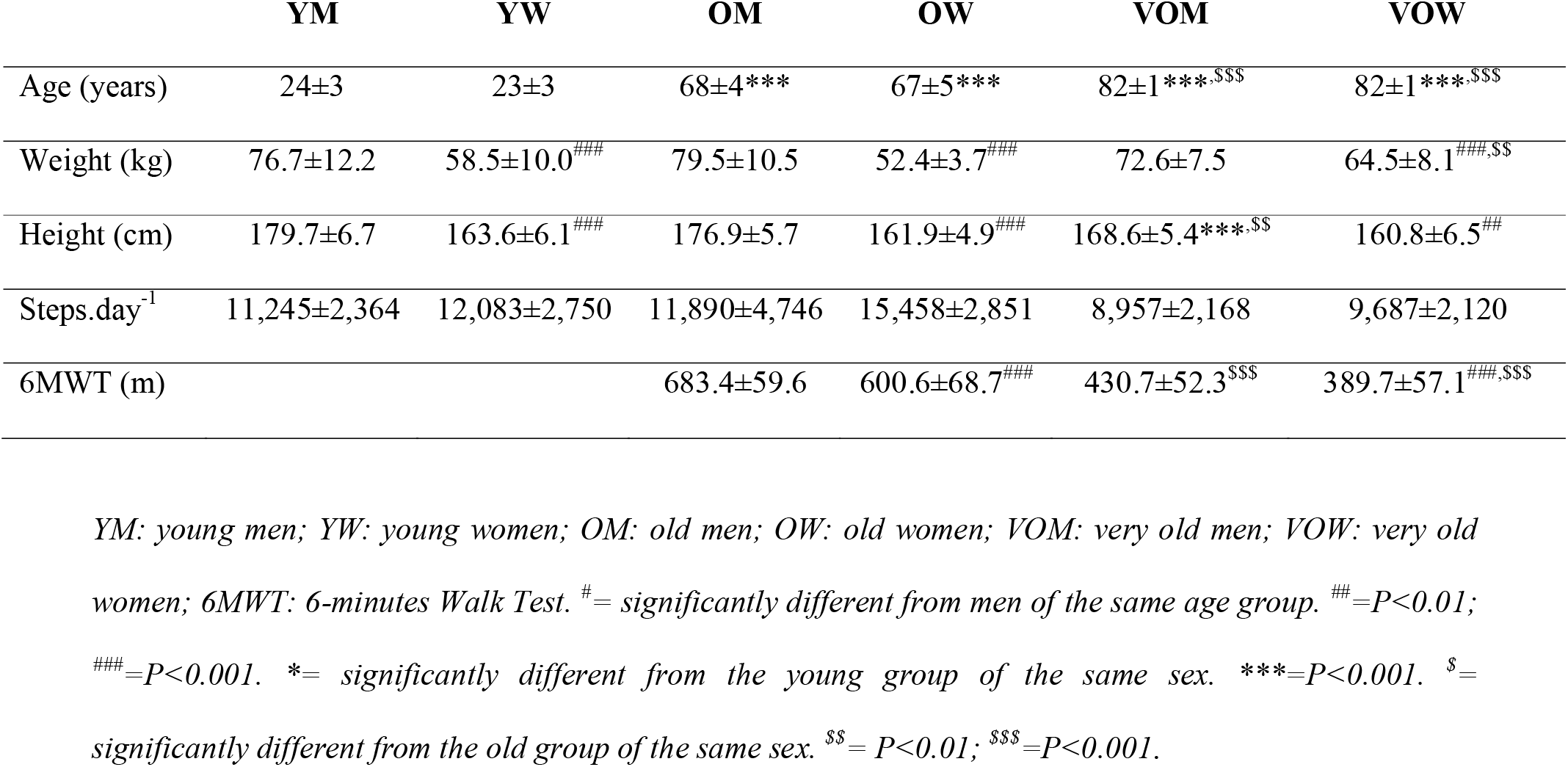
Characteristics of the participants

Men were stronger than women for all age groups. MVIC decreased with age (sex × age interaction: P < 0.001; Table 2). When normalized to body weight, MVIC showed no sex × age interaction (P = 0.545), but decreased with age (age effect, P < 0.001), being greater in men than women (sex effect, P = 0.006; Table 2). Absolute Db_100_ values at baseline showed significant sex × age interaction (P = 0.001). For men, Db_100_ significantly decreased with age (all P < 0.001; Table 2). For women, significant differences were found between YW and VOW, and between OW and VOW (all P < 0.001). No difference was observed between YW and OW (P = 0.407; Table 2). Db_100_ was different between sexes only in the young group (P < 0.001; Table 2). VA did not show main effects of age or sex (P = 0.278 and P = 0.098, respectively), nor age × sex interaction (P = 0.567; Table 2).

**Table 2.**
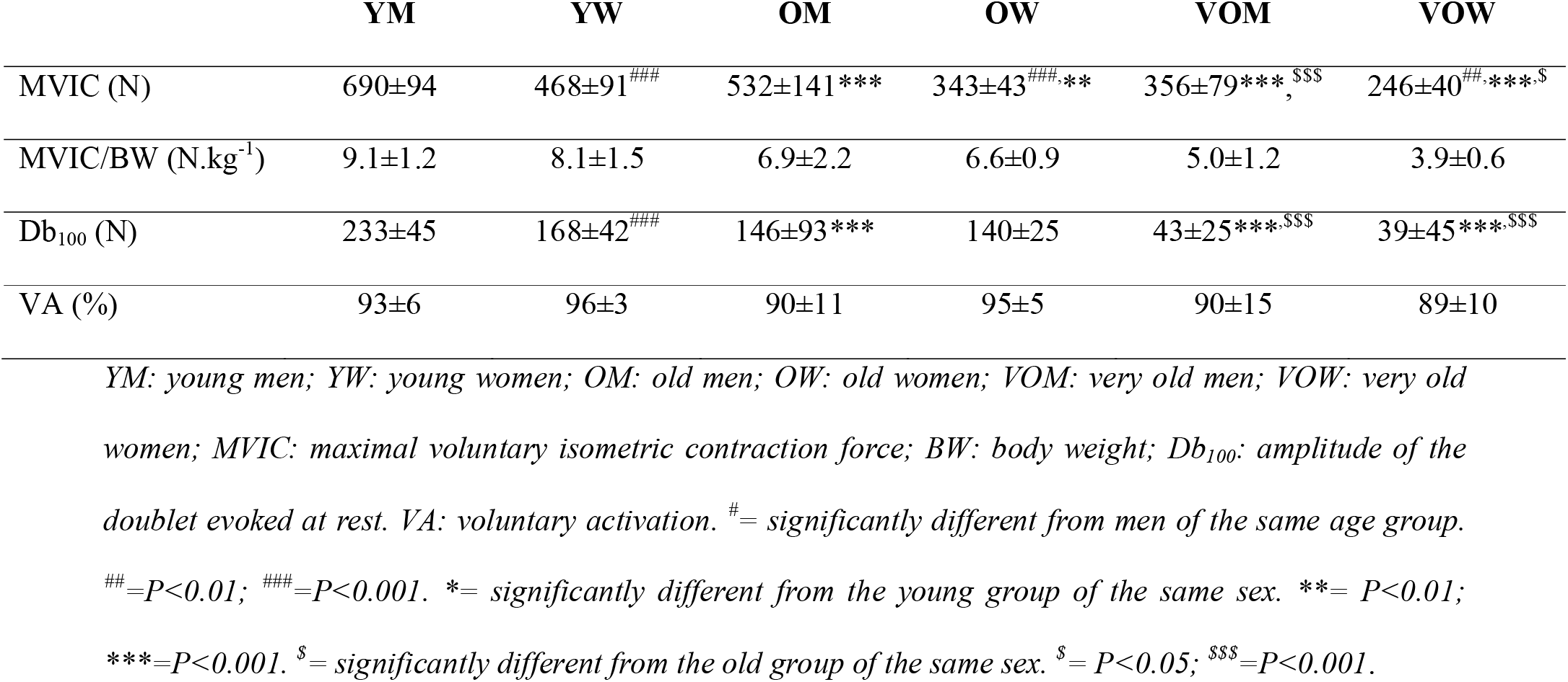
Neuromuscular characteristics of participants at baseline.

### Performance fatigability

Among all the participants, 2 VOM and 4 VOW did not perform the QIF test due to knee pain. Thus, performance data were analyzed for 13 VOM and 11 VOW. Number of contractions was similar across sexes (P = 0.733) but differed with age (P = 0.008). Post-hoc analysis indicated that older adults performed more contractions than young (P = 0.046) and very old adults (P = 0.007), without differences between young and very old adults (P = 1.000, Figure 1A). No significant age × sex interaction was found for number of contractions (P = 0.218). Total work showed significant main effects of age (P < 0.001) and sex (P < 0.001), but no age × sex interaction (P = 0.949). Post-hoc analyses showed greater total work for young and old adults compared to very old adults, without differences between young and old adults, and greater total work for men compared to women (Figure 1B).

**Figure 1.**
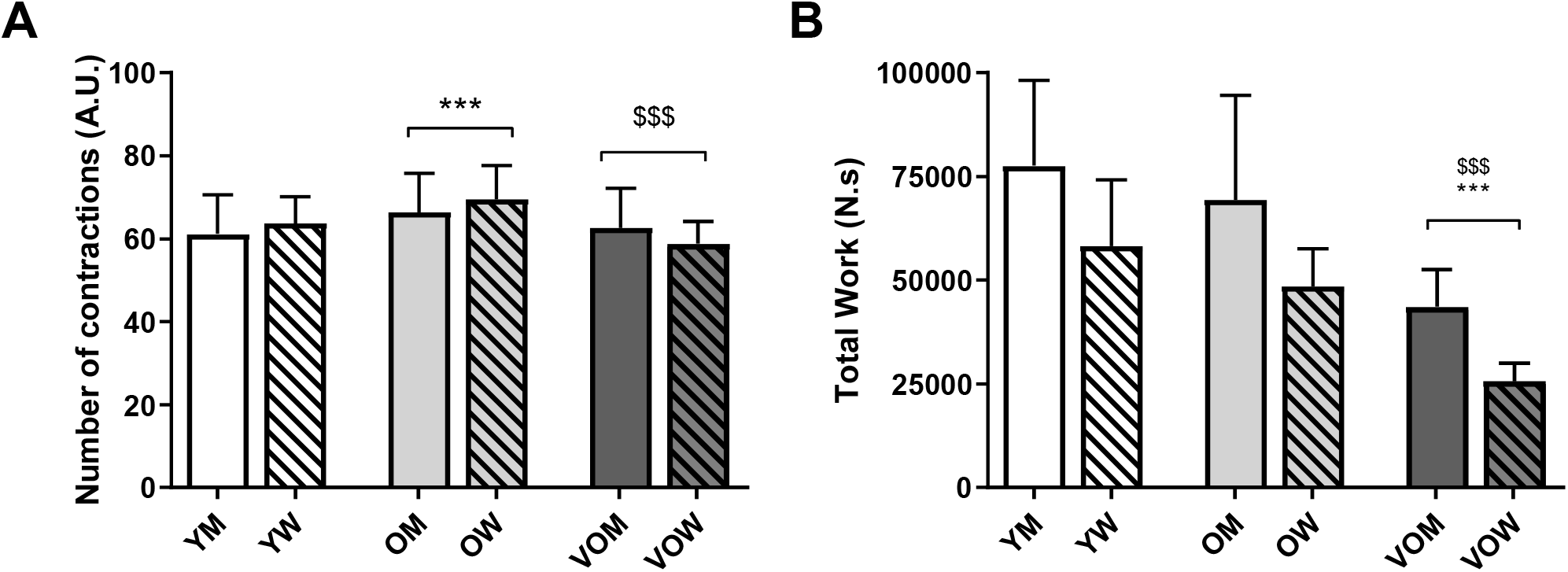
Total number of contractions (panel A) and total work performed at the quadriceps intermittent fatigue test (panel B) for young, old and very old men and women. Total work was greater for men than women in all age groups (all P<0.05). Squared parentheses indicate significant main effect of age. YM: young men; YW: young women; OM: old men; OW: old women; VOM: very old men; VOW: very old women. ***= significantly different from the young group (P<0.001). $$$= significantly different from the old group (P<0.001).

No main effect of sex nor sex × stage, sex × age or sex × stage × age interactions were found for the changes in MVIC, Db_100_ and VA during and after the QIF test (all P > 0.05). Thus, performance fatigability data for men and women were pooled together.

A significant stage × age interaction was observed for the decrease in MVIC relative to baseline (P < 0.001; Figure 2A). The decrease in MVIC was greater for very old adults than young adults at stages 1, 2 and 3. At exhaustion, MVIC loss was greater for young (−28 ± 9%) compared to old adults (−19 ± 8%), but not very old adults (−23 ± 8%; Figure 2A). Db_100_ showed a significant effect of stage (P < 0.001; Figure 2B), indicating that the response to FNMS decreased throughout the test without age-related differences. Significant age × stage interaction was found for VA (P = 0.027; Figure 2C), with a lower voluntary activation at stage 1 for very old adults compared to baseline and young adults.

**Figure 2.**
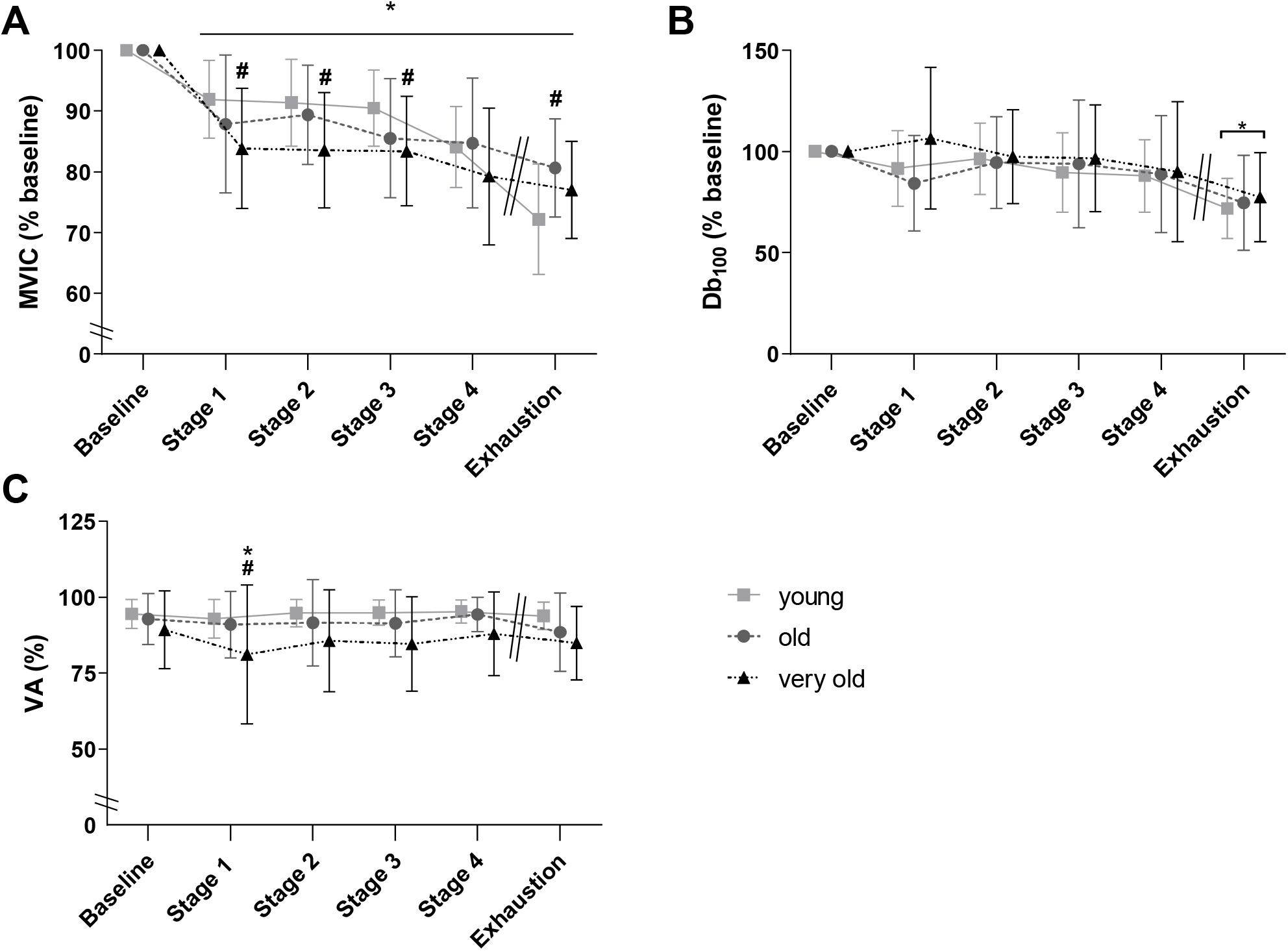
Maximal isometric force (MVIC, Panel A), doublet amplitude (Db_100_, Panel B) and voluntary activation (VA, Panel C) during the QIF test for young, old and very old adults. Because no effect of sex was observed, men and women were pooled together. Squared parenthesis in panel B indicates significant effect of stage. *= significantly different from pre (P < 0.05). ^#^= significantly different from young adults (P < 0.05).

## Discussion

The present study investigated the age-related differences in performance and performance fatigability using a standardized isometric fatiguing test in men and women. Greater relative performance (number of contractions) was observed for old adults compared to young and very old adults, independent of sex. Similar total work was observed for young and old adults, with very old adults showing the lowest total work values. After the QIF test, force loss was greater for young compared to old adults without differences with very old adults, independent of sex. Impairments in contractile function were similar across ages, with no changes in voluntary activation.

### Neuromuscular characteristics at baseline

Maximal force production capacity (absolute and relative to body weight) decreased with age for both sexes. This could be explained by a greater fat mass accumulation with advancing age, lowering the force-body weight ratio, or by a decrease in muscle quality (strength per unit of muscle volume). Other than age, a decreased muscle quality could be induced by the lower physical activity of very old adults compared to the other age groups. Indeed, physical activity has a positive effect on the maintenance of muscle quality with ageing [15], and appears particularly important in maintaining muscle function and autonomy in very old adults [5]. Furthermore, the force evoked at rest by FNMS decreased with age, in agreement with previous studies evaluating the KE muscles [16,17,18,19], evidencing a loss of muscle contractile tissue with age. VA at baseline was similar between age groups. With age, a moderate alteration in VA was previously observed, but results are heterogeneous between studies [20]. The absence of age-related differences in VA in the present study might be due to the relatively high physical activity level of the participants [21], which has been shown to preserve the capacity to activate the working muscles [11,20]. It is possible, however, that the effectiveness of FNMS was limited by the fat thickness of older and very old adults [14], limiting the interpretation of those results despite our precautions.

### Performance and performance fatigability

Old adults performed more contractions compared to the other two groups. However, because young adults were stronger than old adults, total work performed (absolute performance) was similar between those two groups. MVIC loss at exhaustion was greater for young than for old adults, but not very old adults. These results indicate greater fatigue resistance for the old adults (similar absolute performance with reduced force loss), but not very old adults (similar force loss, but lower absolute performance) compared to young adults. In older adults, the preferential loss in fast-twitch motor units is associated with an increase in slow-twitch motor units proportion and size with age, resulting in a relatively higher reliance on the oxidative metabolism [6,22,23,24]. This would lead to a greater fatigue resistance for old than young adults during isometric tasks at relative workloads [6,19,22]. Similar results were previously found on middle-age adults compared to young adults using a similar QIF test [25].

A novel finding of the present study is the loss of greater fatigue resistance in very old adults compared to young adults during isometric tasks of the KE. This agrees with previous evidences on the elbow flexors during an isometric intermittent time to exhaustion task at 60% MVIC [7]. These authors did not observed any differences in time to exhaustion or performance fatigability etiology between young and very old adults, despite the greater absolute workload in young adults [7]. In very old adults, muscle atrophy and loss of both slow- and fast-twitch motor units would decrease neuromuscular function compared to old adults [1,26]. We reported greater force loss at low workloads (*i*.*e*. 10%, 20% and 30% of MVIC) for very old compared to young adults. At exhaustion, MVIC percentage loss of very old adults was between that of young and old adults, without significant difference from the latter two groups, as already reported for concentric tasks [9].

No sex-related differences in performance fatigability were observed for the three age groups. Similar results were reported during intermittent isometric task of the plantar dorsiflexors [27] and concentric task of the KE [9]. Conversely, women showed lower performance fatigability compared to men after a sustained 120-s KE MVIC independent of age [19], probably because they performed lower total work [19,28]. Taken together with the literature, the present results indicate that sex-related differences in performance fatigability are task-specific in young and older adults.

The significant drop in Db_100_ for the three age groups indicated impairments in contractile function throughout the QIF test, without age- or sex-related differences. Different results have been observed in studies comparing young adults with old [6] and very old [7,9] adults, probably because of the different fatiguing protocols adopted [29]. In isometric mode, blood flow occlusion induced by sustained contractions impedes oxygenated blood to reach the capillaries [30]. This facilitates impairments in contractile function [31], attributed to an alteration in excitation-contraction coupling linked with high ATP turnover, free phosphate and H^+^ accumulation [32]. Metabolites accumulation in the working fibers and the recruitment of fast-twitch motor units, composed by type II fibers (less oxygen dependent [30]), could impair performance particularly in very old adults [33]. Regarding central impairments, high levels of physical activity of our participants might prevented an observable alteration in VA, as previously reported comparing trained and untrained individuals [25]. Nevertheless, differences in fatigability in absence of impairments in VA have already been observed in studies investigating sex- [34] and age-related differences [9].

## Conclusions

The present study showed that the greater fatigue resistance of old adults during an isometric intermittent fatiguing task of the KE was not maintained at very old age compared to young adults. Performance fatigability at exhaustion was likely due to impairments in contractile function for the three age groups. Finally, performance fatigability and its etiology were similar between men and women independent of age.

